# Voodoo Machine Learning for Clinical Predictions

**DOI:** 10.1101/059774

**Authors:** Sohrab Saeb, Luca Lonini, Arun Jayaraman, David C. Mohr, Konrad P. Kording

## Abstract

The availability of smartphone and wearable sensor technology is leading to a rapid accumulation of human subject data, and machine learning is emerging as a technique to map that data into clinical predictions. As machine learning algorithms are increasingly used to support clinical decision making, it is important to reliably quantify their prediction accuracy. Cross-validation is the standard approach for evaluating the accuracy of such algorithms; however, several cross-validations methods exist and only some of them are statistically meaningful. Here we compared two popular cross-validation methods: record-wise and subject-wise. Using both a publicly available dataset and a simulation, we found that record-wise cross-validation often massively overestimates the prediction accuracy of the algorithms. We also found that this erroneous method is used by almost half of the retrieved studies that used accelerometers, wearable sensors, or smartphones to predict clinical outcomes. As we move towards an era of machine learning based diagnosis and treatment, using proper methods to evaluate their accuracy is crucial, as erroneous results can mislead both clinicians and data scientists.

## Introduction

Machine learning has evolved as the branch of artificial intelligence (AI) that studies how to solve tasks by learning from examples rather than being explicitly programmed. Machine learning has grown massively over the past decades, with countless applications in technology, marketing, and science [1]. Almost every smartphone nowadays includes speech recognition. Social media and e-commerce websites filter contents and recommend products based on the user’s interests, and scientific data, from astrophysics [2] to neurophysiology [3], are analyzed using machine learning algorithms.

In medicine, a great hope for machine learning is to automatically detect or predict disease states as well as assist doctors in diagnosis, using data collected by phones and wearable sensors. A driving factor is that people carry these devices with them most of the time, and thus a growing amount of daily life data, such as physical activities [4,5], is becoming available. Indeed, an increasing number of studies apply machine learning to the data collected from these devices for clinical prediction purposes. Examples include detecting cardiovascular diseases [6], falls [7], measuring rehabilitation outcomes in stroke and amputees [8,10], monitoring Parkinson’s disease symptoms [11–13], and detecting depression [14,15].

The majority of machine learning algorithms used for clinical predictions are based on the supervised learning approach, which can be summarized in the following steps: first, a set of features is computed from the raw sensor data. These features are typically engineered depending on the specific application; e.g., one feature could be the maximum heart rate in a given time interval. Features are then fed into a machine learning classifier, the parameters of which are adjusted to map each input data point (feature vector or *record*) to its corresponding label, e.g., “healthy”. Once the classifier is trained on enough data, it can be used to perform predictions on new subjects using their features; e.g., do their features predict that they are healthy?

A crucial stage of this process is to assess the prediction accuracy of the trained machine learning algorithm. The standard approach is to use cross-validation (CV) [16], where the data is split into smaller subsets, *folds*, and the classifier is trained on all folds except one, which is the test fold. For example, we can split the dataset into 5 folds, train the classifier on 4 folds, and measure its accuracy on the remaining one. This process is iterated until all combinations of training and test folds have been used, and the final accuracy is obtained by averaging over the individual accuracies on the test folds.

When the goal is to build a model that can generalize to new subjects, the proper form of CV is *subject-wise*: here the training and test folds contain records from different subjects; therefore, the performance of the classifier is measured on a new subject whose data has *not* been used for training (Figure 1A). Nevertheless, many studies employ *record-wise* CV, which randomly splits data into training and test folds regardless of which subjects they belong to. Therefore, records from the same subject are present in both training and test folds (Figure 1B). In this way, the machine learning algorithm can find an association between unique features of a subject (e.g., walking speed) and their clinical state, which automatically improves its prediction accuracy on their test data. As a consequence, the record-wise CV method can significantly overestimate the predicted accuracy of the algorithm.

**Figure 1.**
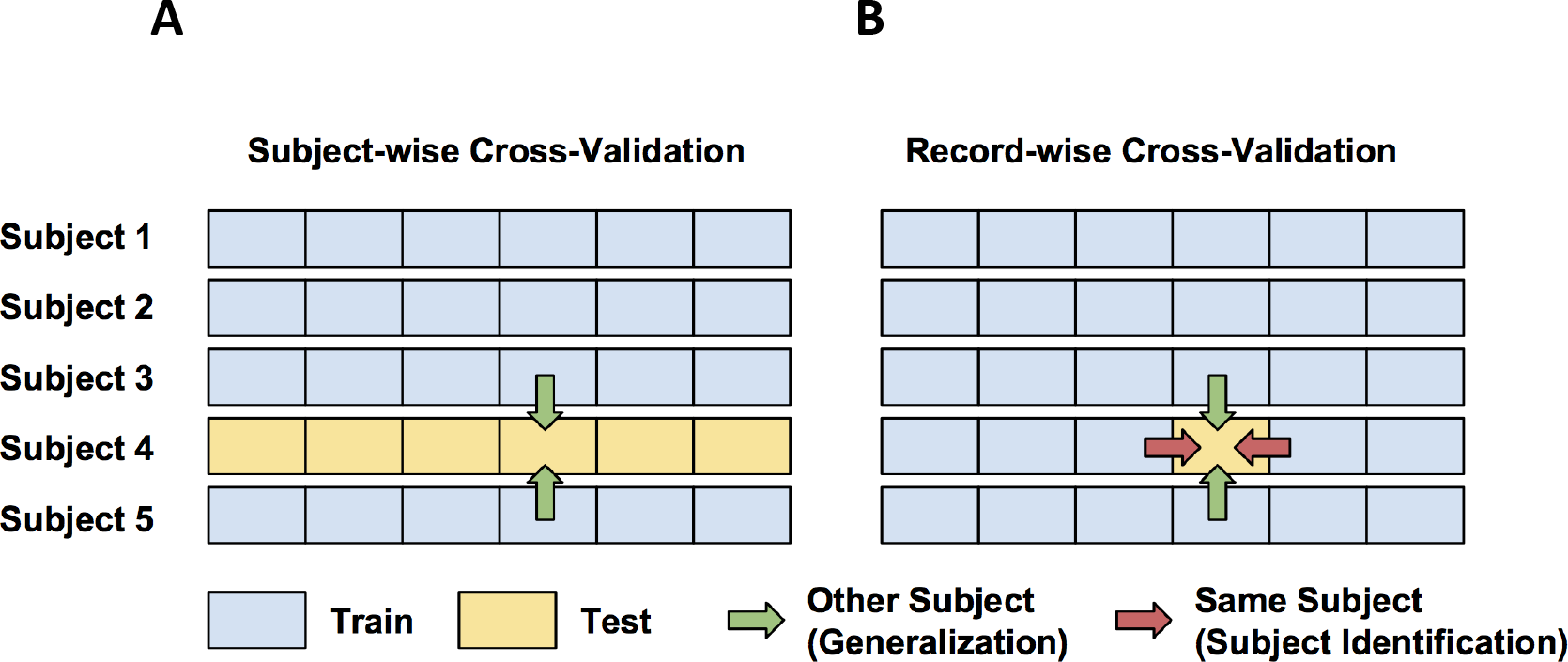
**A visualization of subject-wise (A) and record-wise (B) cross-validation (CV). Data is split into train (blue) and test (yellow) sets to evaluate the performance of machine learning algorithms. Each box represents a record from each subject. While subject-wise CV only uses data from other subjects (green arrows), record-wise in addition uses data from the same test subject (red arrows) to predict its state. The problem is, that if the algorithm can (implicitly) detect the identity of the person based on the features, it can automatically also “diagnose” the disease**.

As this statistical problem is central to our paper, we start explaining the problem with a hypothetical example. Imagine we recruit 4 subjects, 2 healthy and 2 affected by Parkinson’s disease (PD). We have a machine learning algorithm that estimates if a person has PD based on their walking speed. Let our two healthy subjects have constant walking speeds of 1 meter per second (m/s) and 0.4 m/s, and our two PD patients 0.6 m/s and 0.2 m/s. If we do subject-wise CV, we will be unable to predict the performance of the slow healthy subject as well as the fast PD subject, resulting in a prediction accuracy of roughly 50%.Now for record-wise CV, let us have 10 records for each subject. To predict the first of the 10 measurements of the fast healthy subject we would also use the other 9 measurements of that same subject. We would thus be able to certainly conclude that this subject is healthy. The same would be true for the slow healthy subject. After all, none of the PD patients has a walking speed of 0.4 m/s. Subject identification, which we know to be relatively easy, thus replaces disease recognition which we know to be hard. As such, record-wise CV would give us a 100% accuracy which is clearly not supported by the data, and therefore the algorithm will not generalize.

The aim of this paper is (1) to demonstrate the potential threat to the clinical prediction literature when the wrong CV methods are used, and (2) to examine how widespread the problem is. We examine the first aim by showing that record-wise and subject-wise CV yield dramatically different results, with record-wise CV massively overestimating the accuracy. We demonstrate this by using a publicly available dataset on human activity recognition, as well as a simulation that shows how subject-specific and disease-specific factors interact. For the second aim, we present a systematic literature review to quantify the prevalence of this problem in studies that use smartphone-and wearable-based machine learning for clinical predictions. We report the proportion of papers using record-wise CV against those using subject-wise CV, along with their classification accuracies and number of citations.

## Methods

### Activity Recognition Dataset

First, we evaluated how record-wise and subject-wise cross-validation (CV) would be different in a real dataset. We chose a publicly available activity recognition dataset [17] which contained recordings of 30 subjects performing 6 activities: sitting, standing, walking, stair climbing up/down, and laying down. Data consisted of recordings from the accelerometer and gyroscope sensors of a smartphone carried by the subjects. Each data record was a vector of 561 features computed from the sensors signal over a time window of 2.56 s. The dataset contained a total of 10299 records, with an average of 343 records per subject and an approximately equal number of records per activity.

We used random forests [18] for classification. A random forest is an ensemble of decision trees, with each tree providing a prediction about the class of the input data. The forest’s prediction is determined by averaging over the predictions of individual trees. Each tree in a random forest only sees a subset of features and a subset of input data samples. A random forest, thus, has fewer parameters to tune, which makes it less prone to overfitting and a better candidate for generalization to unseen data. We also found random forests to perform well in our previous activity recognition study [19]. Therefore, random forests were an appropriate choice for this study.

For each of the record-wise and subject-wise methods, we used 2, 10, or 30 subjects, and 2, 10, or 30 CV folds. For record-wise, data was randomly split into these folds regardless of which subject it came from. For subject-wise, we split data by subjects such that training and test folds contained records from different subjects. In both methods, the classifier was trained on all but one fold and tested on the remaining fold. The number of trees for the random forest classifier was set to 50, which was optimized based on the out of bag error [18]. We repeated the training procedure 100 times, such that new subjects and folds were randomly generated in each repetition.

### Simulated Dataset

We used a simulated dataset to find out which properties of human subject data make the performance of subject-wise and record-wise CV methods different. Specifically, we were interested in two properties: *cross-subject variability*, and *within-subject variability*. Cross-subject variability is the variability in data that is observed across the subjects. In a diagnosis scenario, this property captures the effect of the disease, or the features that distinguish healthy subjects from sick. Within-subject variability represents the variability observed when multiple samples are recorded from a single subject. In clinical data, this property is usually related to the changes in the disease trajectory, as well as changes in the physiology of the subjects.

We used a generative model to create the simulated dataset. This model combined cross-subject and within-subject variabilities to generate an observed record **y** as the following:
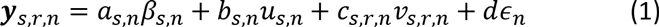
where each record, **y**, is a 3–dimensional matrix, with *s*, *r*, and *n* indicating subject, record, and feature dimensions, respectively. β is an *S⨯N(S*: number of subjects, *N*: number of features) matrix of disease effects, or fixed effects, with rows of 1 for patients and −1 for healthy subjects. ***U***~*N*_S⨯N_(0,1) is an *S⨯N* matrix accounting for the random cross-subject variability, or random effects, ***V***~*N*_S⨯N⨯_(0,1)(*R*: number of records) is an *S⨯N⨯R* matrix representing the within-subject variability, and ɛ~*N*_*N*_(0,1) is the feature-generating *N*-dimensional random vector. Finally, *a*_*s,n*_, *b*_*s,n*_, *c*_*s,r,n*_ and *d* are constant parameters. To simplify the model, we considered *a*_*s,n*_=*a*, *b*_*s,n*_=*b* and *c*_*s,r,n*_=*c* to be constants, where *b* and *c* control the proportions of cross-and within-subject variability, respectively.

Similar to the activity recognition dataset, we trained and tested random forests on the simulated dataset using record-wise and subject-wise methods. For record-wise, we randomly split the generated dataset into 50% training and 50% test, regardless of the subjects. For subject-wise, we did the same but we ensured that same subjects were not present in both training and test sets. We generated datasets with variable number of subjects, from 4 to 32. These numbers were within the range used in the reviewed studies, as well as in the activity recognition dataset. We used *N* = 10 features and *R* = 100 records for each subject. We set *a* = 0.5, accounting for a disease effect size of (1−(−1)) ⨯ 0.5 = 1, and *d* = 0. 1. For each CV method and each value of *b* and *c*, we measured the classification error on the test set. Each simulation was repeated 10 times and the average classification error across the repetitions was calculated.

### Literature Review

We reviewed the literature to find out what percentage of the studies used record-wise versus subject-wise CV. We specifically looked for the studies which used both machine learning and smartphone or wearable sensor technology for clinical predictions. This process had three steps: (1) finding the relevant studies; (2) determining the CV method used by each study; and (3) finding the reported classification error in each study. The papers were reviewed and analyzed by two authors (SS, LL). A total of four discrepancies were resolved by consensus.

**1)*Finding Relevant Studies***.We used Google Scholar [20], and searched for the studies which contained at least one of the following keywords in their title: “wearable”, “smartphone”, or “accelerometer”. In addition, to account for the clinical applicability, the papers had to have one of the following keywords in their text: “diagnosis”, “disease”, or “rehabilitation”. Finally, the papers had to contain “cross-validation” at least once, and be published after 2010. These criteria resulted in the following Google Scholar search query:

(*intitle:wearable OR intitle:smartphone OR intitle:smartphones OR intitle:accelerometer) AND (diagnosis OR disease OR rehabilitation) “Cross-Validation” (from 2010*)

From the results returned by this query, we excluded papers which met at least one of the following conditions:

1. used one data point (record) for each subject. In fact, when there is one sample per subject, record-wise CV and subject-wise CV are the same.
2. used *personal* models, which are trained and tested on each individual subject separately.
3. did not use any CV procedure.
4. did not relate to any clinical or rehabilitation application.
5. were review of the literature.
6. were not peer-reviewed (e.g., reports).
7. did not allow us to determine the CV type (see Determining CV Type).

**2)*Determining CV Type***.After finding the relevant papers, we grouped them into the ones which used record-wise CV and the ones that used subject-wise CV We assigned a paper to the subject-wise group if one or more of the following conditions were satisfied:

1. The authors used the term “subject-wise” when explaining their cross-validation strategy.
2. The authors used the term “Leave-One-Subject-Out”.
3. The authors mentioned that they tested their algorithms on test subjects which were not included in the training set
4. We did not find any overlap between the subject IDs in their training and test datatsets, where subject IDs were provided.

We assigned a paper to the record-wise group if one or more of the following conditions were satisfied:

1. The number of cross-validation folds was greater than the number of subjects.
2. The authors mentioned that they randomly split the whole dataset into training and test sets.
3. The authors mentioned that they randomly assigned data samples to cross-validation folds.
4. We found an overlap between subject IDs between the training and test datasets, where subject IDs were provided.

**3)*Finding Reported Classification Error***. We also wanted to see if subject-wise and record-wise CV studies were different in their reported classification error (1 - accuracy). Since papers used different metrics to report the classification accuracy of their algorithms, we used the following rules to find their accuracies:

1. Where a single accuracy or classification error was reported, we directly used that.
2. Where multiple accuracies were reported for different conditions or classes, we used the average accuracy.
3. Where accuracies were reported for different types of classifiers or feature sets, we reported the highest accuracy.
4. Where sensitivity and specificity were reported and their difference was less than 2%, we used their average as accuracy. This is supported by the fact that accuracy is bounded by sensitivity and specificity [^*^ By definition, sensitivity = TP/(TP+FN), specificity = TN/(TN+FP), and accuracy = (TP+TN)/(TP+FN+TP+TN), where TP, FN, TN, and FP stand for true positive, false negative, true negative, and false positive, respectively. Since in]
5. Where Fl-score was reported, or precision and recall were reported from which we could calculate the F1-score, we used it instead of the accuracy.
6. We excluded papers which only reported root-mean-square error (RMSE), normalized RMSE (NRMSE) or similar metrics as they are related to estimating a continuous variable and cannot be converted to classification errors.

## Results

### Activity Recognition Dataset

We started by evaluating the performance of subject-wise CV on the UCI activity recognition dataset. When using 2 folds and 2 subjects only, the error rate started at a value of 27% and, as the number of subjects increased to 30, it decreased significantly and reached 9% (Figure 2). Similarly, as the number of folds increased, i.e., data from more subjects was used for training the classifier, the error rate decreased and leveled around 7% with 30 folds (Figure 2, green lines).

We then trained the classifier using record-wise CV and used the same procedure to assess how the error changed as a function of number of subjects and folds. Interestingly, the classification error already started at a value of 2% when using data from 2 subjects, and did not significantly change when either number of subjects or folds increased (Figure 2, orange lines). Therefore, regardless of the amount of data used for training the classifier, record-wise CV significantly overestimated the classification accuracy on this dataset.

**Figure 2.**
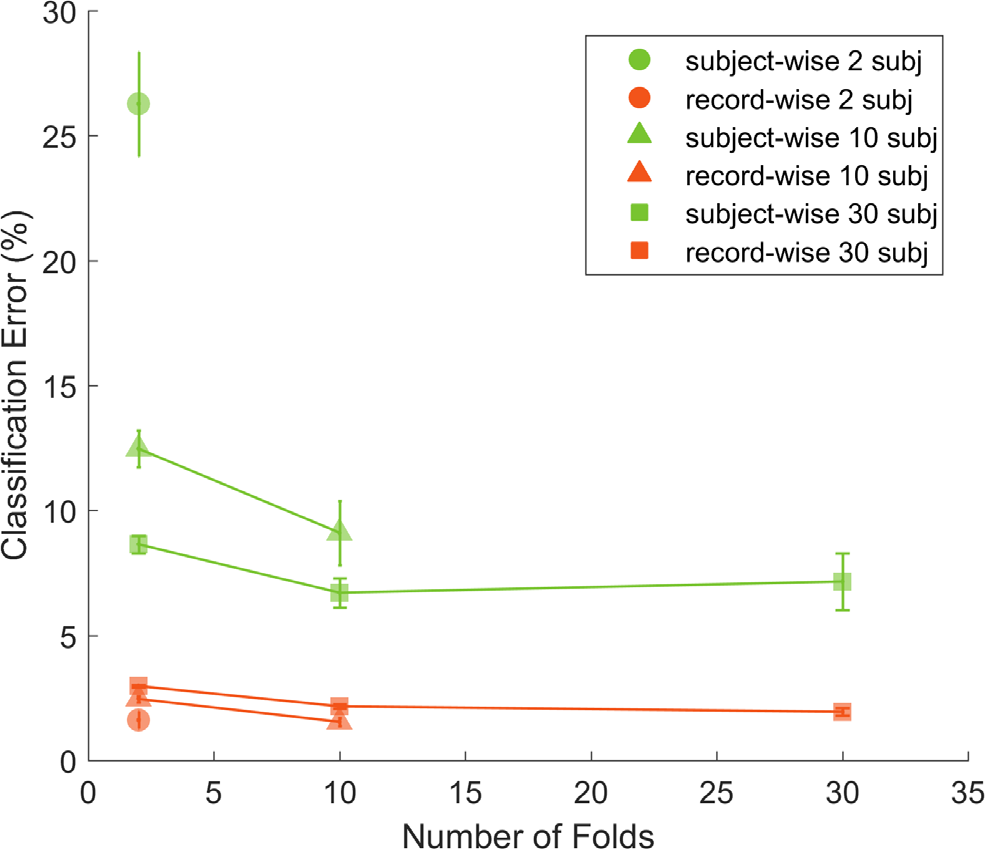
**Effect of Subject-wise and record-wise CV on the classification error for the UCI activity recognition dataset. As the number of folds (x-axis) used to perform CV increases, the error tends to decrease. Similarly, performance improves when the number of subjects increases (symbols denote total number of subjects used to train and test the classifier). Record-wise CV significantly inflates the predicted accuracy (orange) as compared to subject-wise CV (green). Error bars indicate 95% confidence intervals**.

General min(a/b, c/d) ≤ (a+b)/(c+d) ≤ max(a/b, c/d) is true, we can conclude that accuracy is bounded by sensitivity and specificity.

### Simulated Dataset

We quantified how cross-subject variability (*b*) and within-subject variability (*c*) contributed to the classification errors in either of the two CV methods (see Equation 1). As in the activity recognition task, we used a random forest classifier. Figure 3A shows how classification error changed with *b* and *c*, for 4,12, and 32 subjects. In all scenarios, for small values of *b* and *c* the classification error was small. This is because when both within and cross-subject variabilities are low, the disease effect is more prominent strongly present in the features, helping the classifier to distinguish between healthy and diseased subjects. For higher values of *c*, classification errors were higher, especially for the subject-wise method.

**Figure 3.**
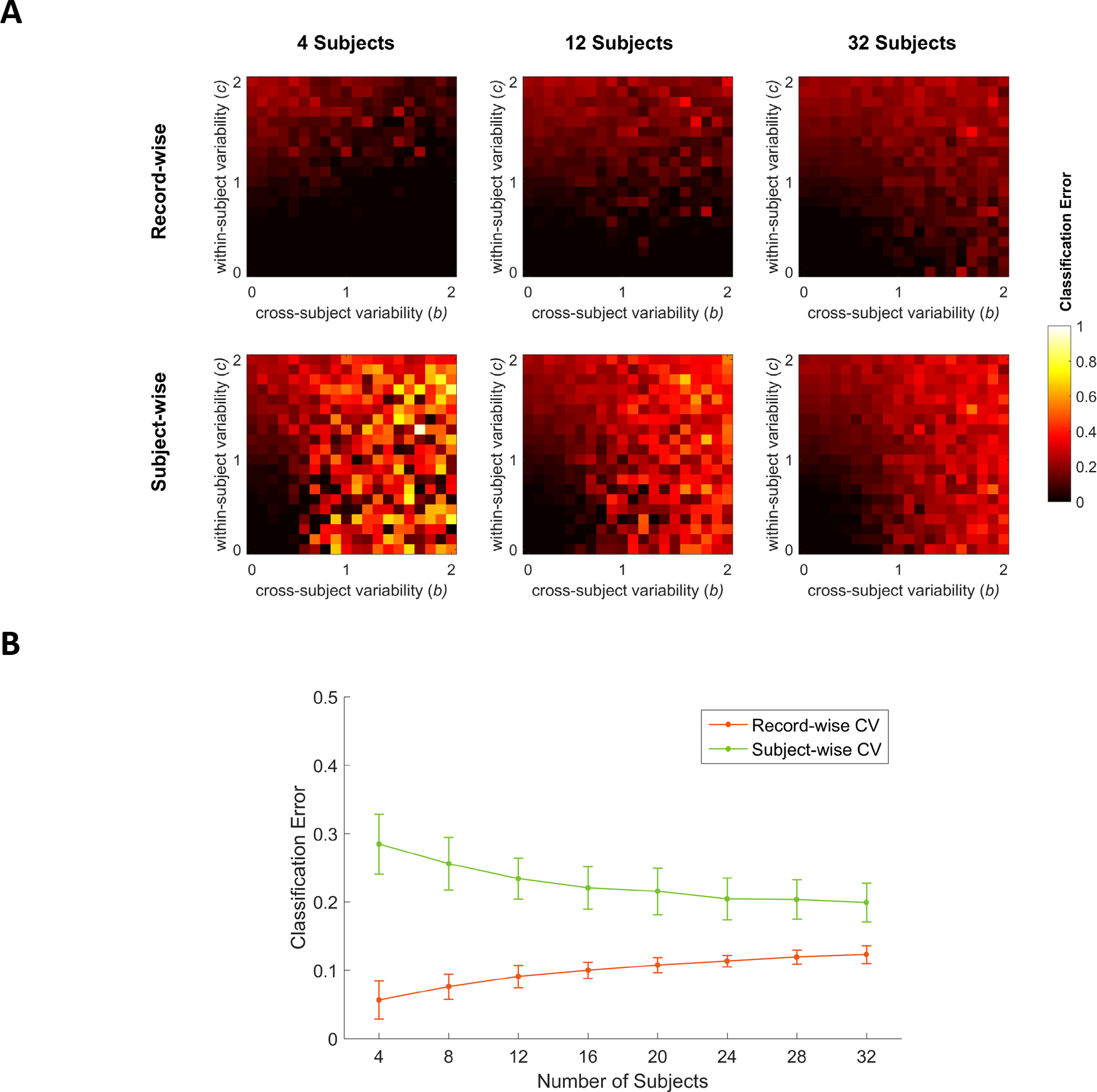
**Classification error in a simulated dataset for record-wise and subject-wise cross-validation (CV) methods, as a function of cross-subject (*b*) and within-subject (*c*) variability (A), and number of subjects (B). The number of features is set to 10. (A) Each column shows the classification error for both CV methods as a function of *b* and *c*. Brighter colors indicate higher classification error values (black = 0; white = 1). (B) The mean and the standard deviation of classification error for subject-wise and record-wise methods, as a function of number of subjects**.

Increasing the cross-subject variability (*b*) alone did not increase the classification error for the record-wise method, when the number of subjects was small (top left panel). Indeed, in the record-wise CV, the classifier was already informed about *b* by having samples from all or most of the subjects in its training set. For the subject-wise method, on the other hand, increasing *b* dramatically increased the classification error (bottom left panel). Nevertheless, when more subjects were used, the classification error increased in both CV methods, but remained lower for the record-wise method (top right versus bottom right panel).

Overall, as shown in Figure 3B, record-wise CV underestimated the classification error, relative to the subject-wise CV method. This difference was largest when the number of subjects was small, and gradually decreased as we increased the number of subjects. This is because as we increase the number of subjects, it becomes harder for the record-wise classifier to identify subjects based on their data, thereby losing its advantage over the subject-wise method. Nevertheless, even for relatively large number of subjects, record-wise CV leads to a significantly lower classification error than subject-wise CV.

### Literature Review

Finally, we wanted to understand how frequently either of the subject-wise and record-wise CV methods was used in the relevant literature. Our search query returned a total of 369 papers, of which we analyzed the first 200. Out of these, 55 papers matched our inclusion criteria, with 24 (43.6%) using record-wise CV and 31 (56.4%) using subject-wise CV. Figure 4 summarizes the literature review procedure.

**Figure 4.**
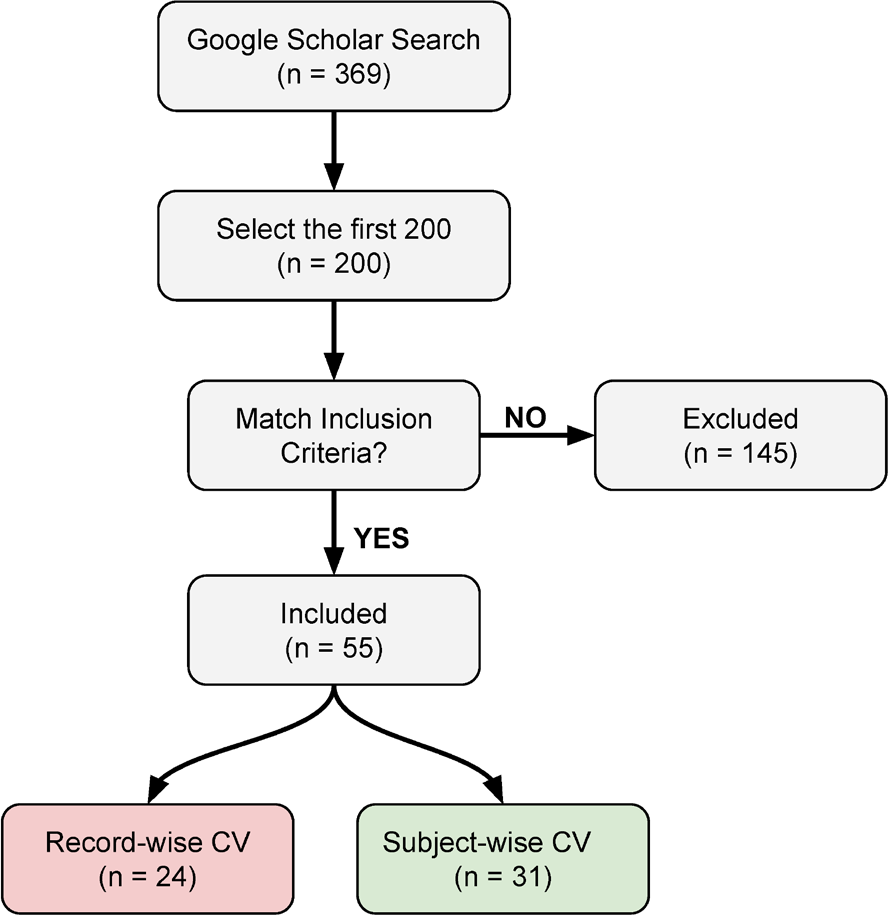
**Flowchart diagram of the literature review procedure**.

Then, we extracted the reported classification errors from the papers included in our study. Due to lack of information, we could only extract this from 42 out of the 55 papers. As shown in Figure 5A, for subject-wise CV papers, the median classification error was 14.15%, more than twice that of record-wise CV, which was 5.80%. These values were significantly different (P<0.01, two-tailed Wilcoxon rank sum test), exhibiting an inflated performance yielded by record-wise CV. Therefore, improper crossvalidation has led to dramatically “better” results.

Interestingly, the median number of citations received by papers in each category were very close as shown in Figure 5B, with subject-wise studies receiving 10 and record-wise 9 citations per paper. Therefore, whether a paper used the correct or wrong CV method did not affect the perceived impact of that paper in the field.

**Figure 5.**
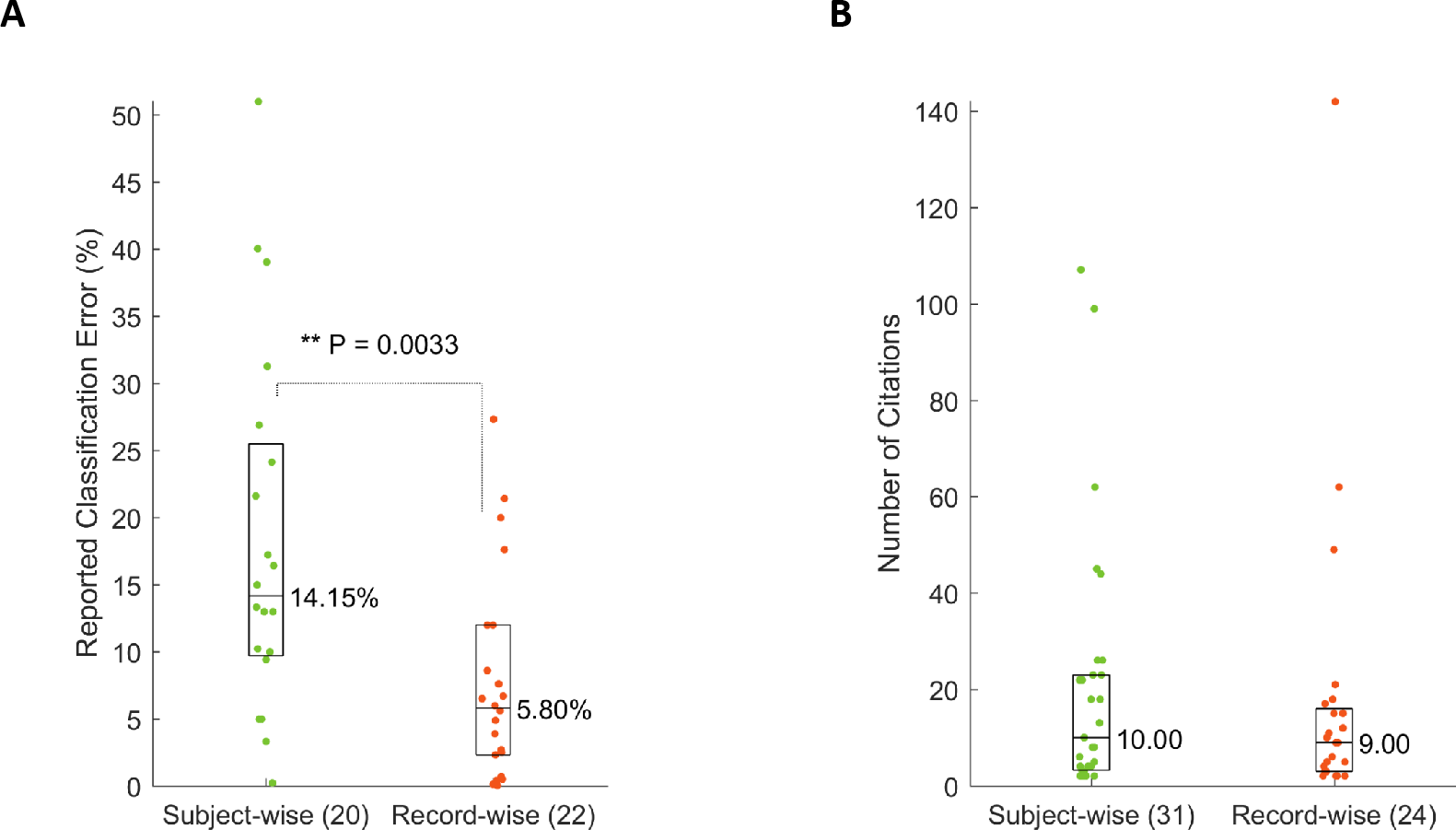
**Literature review results. Each plot, shows the distribution of the variable of interest for subject-wise CV (SWCV) and record-wise CV (RWCV) methods. Boxes show the interquartile range, and the horizontal lines inside the boxes indicate the medians. A: reported classification error, inferred as detailed in Literature Review. B: total number of citations each paper has received**.

## Discussion

We evaluated the reliability of reported accuracies in studies that used machine learning and wearable sensor technology to predict clinical outcomes. Using a publicly available dataset and a simulation, we first showed that the record-wise CV method produces misleadingly high accuracy. Then, we performed a literature review and found that about 44% of the studies used record-wise CV. As expected, the accuracies reported by the studies using the wrong method (record-wise) were higher than the ones reported by correct (subject-wise) papers. Therefore, it seems that a significant proportion of studies are dramatically overstating the ability of current machine learning algorithms to predict clinical outcomes.

Here we only considered one way that cross-validation can go wrong. In fact, the problem is not limited to using the record-wise method. In many prediction algorithms, there are hyperparameters that need to be adjusted, and very often, these are chosen such that the prediction error is minimized. This makes the algorithms overfit the data and thus not generalize to other datasets. The correct way for learning these hyperparameters is to further divide the training set into training and validation subsets, and then minimize the prediction error on those validation subsets rather than on the whole dataset. This approach allows for a proper evaluation of the generalizability of the algorithm [16].

Furthermore, while we only used one dataset, one could use any clinical dataset to show how mixing the subjects between training and validation sets can artificially increase the prediction accuracies. Clinical records often include information about physical or physiological characteristics of an individual, such as waist size or blood type, which do not vary much over time. For such data, if we use record-wise CV, the algorithms will already know the clinical outcome for an individual with specific characteristics. On the other hand, subject-wise CV will ensure that there is no way for the algorithm to exploit such shortcuts. Therefore, choosing the right CV method is particularly important in clinical prediction applications.

The use of machine learning for clinical predictions is growing in popularity [21]. This is in part because the computing power of electronic devices has increased to the point that current mobile devices are as powerful as supercomputers from a few decades ago. This means that we are now able to run very complex algorithms, on enormous amounts of data, using relatively cheap devices in a short period of time. In addition, the emergence of more advanced measurement devices with high spatial and temporal resolution (e.g., [22,23]) requires the use of techniques that can analyze such large amounts of high-dimensional data, and in those applications, machine learning is replacing traditional data analysis. As such, machine learning tools are now used by investigators who might not have proper training, which can open the door to misuse.

The most advanced algorithms and the highest quality datasets, if used with wrong CV methods, lead to meaningless results. Such erroneously positive results threaten the progress in the field. A direct consequence is the waste of resources on the projects that are based on fallacious findings.

Furthermore, such results might contribute to the problem of irreproducibility of research findings [24,25] and thereby undermine the trust in both medicine and data science. Only with meaningful validation procedures can the transition into machine learning driven, data-rich medicine succeed.

## Acknowledgements

This study was supported by the following National Institute of Health grants: 5R01NS063399, P20MH090318, and R01MH100482. Authors AJ and LL were supported by CBrace 80795 Otto Bock Healthcare Products, GmBH.

